# A Hit Prioritization Strategy for Compound Library Screening Using LiP-MS and Molecular Dynamics Simulations Applied to KRas G12D Inhibitors

**DOI:** 10.1101/2025.04.22.650081

**Authors:** Foroughsadat Absar, Brandon Novy, Edith Nagy, Evgeniy V. Petrotchenko, Konstantin I. Popov, Jason B. Cross, Roopa Thapar, Christoph H. Borchers

**Affiliations:** Segal Cancer Proteomics Centre, Lady Davis Institute, Jewish General Hospital, McGill University, Montreal, Quebec, H3T 1E2, Canada; Division of Chemical Biology and Medicinal Chemistry, University of North Carolina at Chapel Hill, Chapel Hill, NC 27599, USA; Institute for Applied Cancer Science, The University of Texas MD Anderson Cancer Center, Houston, TX 77030, USA; Gerald Bronfman Department of Oncology, Jewish General Hospital, McGill University, Montreal, Quebec, H3T 1E2, Canada; Division of Experimental Medicine, McGill University, Montreal, Quebec, H3A 1A1, Canada; Department of Pathology, McGill University, Montreal, Quebec, H3A 1A1, Canada

## Abstract

An important step in screening small molecule libraries for drug discovery is hit prioritization and validation to rule out false positives, which is usually performed using biochemical and biophysical assays. The development of orthogonal assays that are highly sensitive and can accelerate the hit-to-lead process is valuable. Limited proteolysis combined with mass spectrometry (LiP-MS) is a technique used to study changes in protein structure upon ligand binding. In LiP-MS, proteins are exposed to low concentrations of proteases under native conditions. The resulting proteolytic pattern is sensitive to protein structure at the cleavage site, which can change upon ligand binding. We characterized the interaction of small molecule inhibitors of the KRas G12D mutant oncoprotein by LiP-MS combined with molecular dynamics (MD). Intact mass spectrometry and top-down analysis were used to detect and identify KRas G12D cleavage products in the presence and absence of inhibitors, thereby locating the cleavage sites in the protein. Cleavage sites protected upon compound binding correlated well with the switch II binding site. The degree of cleavage depends on binding affinity and the presence of specific functional groups in the inhibitor’s structure. A comparison of MD simulations for the ligand-free and ligand-bound proteins revealed the atomistic mechanisms by which the cleavage sites, located in flexible and disordered regions, are stabilized upon compound binding. We suggest that LiP-MS combined with MD (LiP-MS-MD) could be valuable in small molecule screening campaigns and add to the repertoire of available methods for high-quality hit selection in early-stage drug discovery.

## Introduction

Limited proteolysis combined with mass spectrometry (LiP-MS) is an informative technique for studying changes in protein structure (1). In this method, a non-specific proteolytic enzyme is used to cleave the protein for a short time. The first cleavage event occurs while the protein’s tertiary and quaternary structures are preserved. This method has a wide range of applications due to its ability to report on protein structure, dynamics and interaction with other ligands under different conditions. It has been used to study protein conformational changes, protein small-molecule binding, and protein-protein interactions (2-9). This makes LiP-MS particularly valuable in fields such as functional and structural proteomics as well as drug discovery, where it helps identify binding sites and elucidate the roles of target proteins (3, 4, 9-11).

Here, we have used LiP-MS to characterize the mode of binding of five non-covalent pyrido[4,3-d]pyrimidine inhibitors that were recently developed and are specific to the KRas G12D oncoprotein (12). We quantitatively compared a single cleavage product in free and compound-bound protein states, making the LiP-MS part of the method faster, more robust and useful for screening hit prioritization.

The Rat sarcoma viral oncogene (Ras) family of proteins is a group of GTP/GDP molecular switches and the American Cancer Society estimates that they are frequently mutated in ∼19% of all human cancers (13). The KRas G12D mutation is a highly prevalent oncogenic driver and is frequently found in pancreatic ductal adenocarcinomas (PDAC) (44%) where it is associated with low survival rates and high mortality (14-16). It is also present in approximately 10-12% of colorectal cancer (CRC) cases, leading to resistance to anti-EGFR therapy and a poor prognosis (17, 18). KRas G12D is less common in non-small cell lung cancer (4%), but it is linked to immunosuppression and resistance to the immune checkpoint inhibitors (19-21). The high prevalence and poor prognosis associated with KRas G12D mutations in various cancers make it an attractive therapeutic target. Recent advances in targeting KRas G12D have introduced promising strategies to inhibit this oncogenic driver. Approaches such as the NS1 monobody disrupt the α4-α5 interface of KRas, thereby impairing signaling and tumor growth (22, 23). Small molecule inhibitors, including thieno[2,3-d]pyrimidine derivatives, have demonstrated selective reduction of KRas G12D activity and downstream signaling in cancer models (24). Salt bridge formation with the Asp12 residue offers another precise mechanism for binding and disrupting KRas interactions (25). Furthermore, the development of RMC-9805, a first-in-class, oral, mutant-selective covalent inhibitor, has shown promising efficacy in preclinical models by forming a stable tri-complex with cyclophilin A, effectively disrupting KRas G12D signaling and leading to tumor regression (26). Similarly, a novel covalent approach utilizing malolactone-based inhibitors selectively targets KRas G12D through strain-release alkylation, forming stable covalent complexes that inhibit cancer cell proliferation and tumor growth (27). Recently, Mirati Therapeutics disclosed MRTX1133 as a potent and highly selective KRas G12D inhibitor, demonstrating over 1000-fold selectivity for the mutant protein compared to wild-type KRas, with picomolar affinity for KRas G12D and significant antitumor efficacy in preclinical models (12, 28). This inhibitor has entered Phase I/II clinical trials (29). Several compounds belonging to the MRTX1133 pyrido[4,3-d]pyrimidine scaffold that bind KRas G12D with a range of affinities have also been disclosed (US Patents WO2021041671, WO2022031678, WO2022132200).

Designing ligands that modulate protein function by stabilizing specific conformations is a widely used strategy in drug discovery (30). This approach is particularly relevant for targeting proteins with conformational flexibility essential to their function (31). It was found that variations in Ras dynamics can be explored in developing isoform- and mutant-specific small molecule inhibitors (32). For example, a distinct conformation of KRas(G12D) is pivotal for the design of a mutant-selective of monobody inhibitors (33). Additionally, [(2R)-2-(N’-(1H-indole-3-carbonyl)hydrazino)-2-phenyl-acetamide] was found to be a conformationally specific small molecule binder inhibiting KRas-mediated signal transduction for KRas mutants (33, 34). These examples highlight the need for a hit-to-lead optimization pipeline that not only focuses on protein-ligand binding affinity but also characterizes ligands based on their ability to stabilize protein structure upon target engagement.

Given the importance of KRas G12D in cancer, the availability of well-characterized chemical matter for this target, and the need for a structure-informed approach, we assessed the potential of using LiP-MS in proof-of-concept experiments for the KRas G12D protein. We used intact protein mass spectrometry to detect and top-down analysis to identify KRas cleavage products, thus locating the cleavage sites in the protein sequence. Inhibitor binding stabilized the protein structure reducing the degree of cleavage. The extent of the proteolysis reaction was measured quantitatively using intact protein mass spectrometry and was found to be dependent on the ligand binding affinity and certain functional groups in the inhibitor scaffold.

To complement the results obtained by LiP-MS, we conducted molecular dynamics simulations alongside molecular docking to provide atomistic insights into the protein-ligand interactions and the mechanism of ligand-induced local protein stabilization. The combination of proteomics techniques and molecular dynamics simulations has previously been used to study a number of biological problems, such as protein structure determination (35), metal-binding events (36), and lipid-protein interactions (37, 38, 39). However, to our knowledge, this study represents the first integration of LiP-MS and molecular dynamics to investigate the effects of ligand binding on protein dynamics. Our results show that this approach can be used for the identification of the inhibitor binding sites, characterization of protein-ligand interactions and potentially for secondary screening of small-molecules libraries.

## Results and Discussion

We used LiP-MS analysis to characterize the binding of KRas G12D protein with five inhibitors of differing structures and binding affinities. We observed significant protection from proteolysis upon inhibitor binding with varying degrees of cleavage for different compounds. The obtained results were explained by comparing MD simulations of free and compound-bound protein-ligand complexes, providing an atomic-level description of the conformational changes upon ligand binding. These simulations revealed key intermolecular interactions and local structural fluctuations in the KRas G12D mutant, that are likely responsible for the observed proteolytic patterns.

### Detection of protection from proteolysis of KRas G12D protein upon inhibitor binding

The proteolytic pattern of the KRas G12D protein showed significant changes upon compound binding, as was observed in SDS-PAGE analysis. The analysis indicated that binding of Compound 5 led to significant protection against enzymatic cleavage by both proteinase K and chymotrypsin (Figure 1A, 1B). In the free state, KRas G12D was extensively cleaved into smaller peptides, while in the compound-bound state, the protein displayed a distinct reduction in cleavage, suggesting stabilization of the protein structure upon ligand binding. The same finding was confirmed by intact protein mass LC-MS analysis (Figure 1C, S1). We have observed the appearance of new protein charge envelopes corresponding to the cleavage products of the KRas G12D protein for both proteinase K and chymotrypsin enzymes.

**Figure 1.**
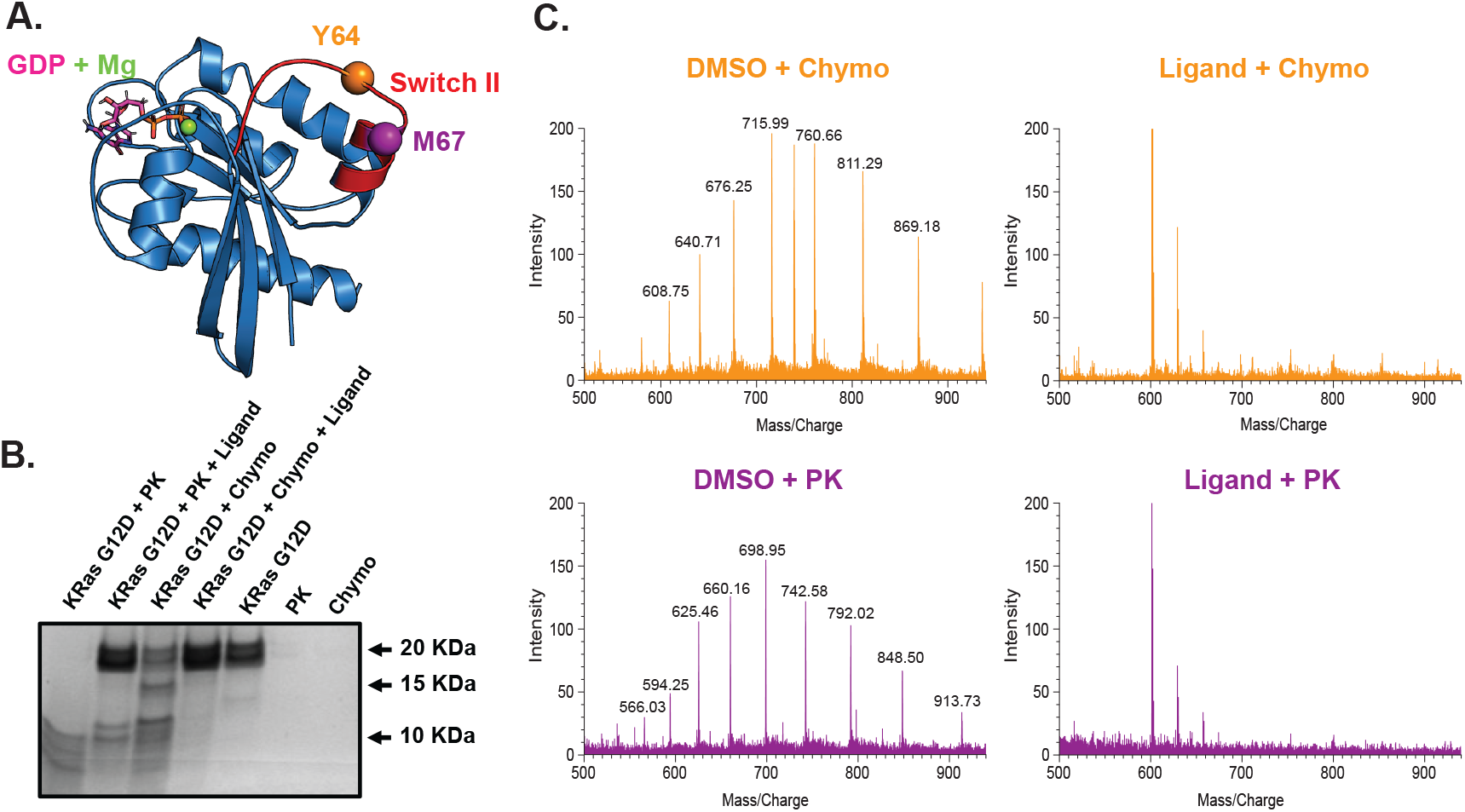
Limited proteolysis of KRas G12D in free and compound-bound states. **A**. Crystal structure of the KRas G12D – 6IC ligand complex (PDB: 7RPZ). The identified cleavage sites are located in the switch II loop (red), the GDP and Mg cofactors are highlighted in pink and green, respectively. **B**. SDS-PAGE analysis of limited proteolysis reactions of KRas G12D with and without Compound 5 (MRTX1133) using two proteolytic enzymes (proteinase K and chymotrypsin). Lane 1, 2 – reaction with proteinase K; lane 3, 4 – reaction with chymotrypsin; lane 5 – intact KRas G12D; lane 6, 7 – proteinase K and chymotrypsin controls. There is significant protection against the KRas G12D cleavage in the presence of Compound 5. **C**. Intact protein LC-MS analysis of KRas G12D limited proteolysis reactions using chymotrypsin (top, orange) and proteinase K (bottom, purple) without (left) and with (right) Compound 5. From the representative charge envelopes displayed, there is a noticeable absence of the protein cleavage products in the ligand-bound samples.

### Identification of the limited proteolysis cleavage products by LC-MS

The LC-MS analysis of the limited proteolysis digestion products allowed for precise mapping of the cleavage sites in the KRas G12D protein. In the absence of the inhibitor, the total ion chromatogram revealed distinct peaks corresponding to specific cleavage products. For proteinase K, a cleavage product with a mass of 11,865.0 Da was identified by matching the mass to all possible KRas G12D peptide masses, pinpointing residue M67 as the cleavage site. This was confirmed by top-down analysis with CID fragmentation using a precursor ion of m/z 792.0^+15^, which identified the product as the C-terminal portion of the protein (aa68-aa169). Similarly, for chymotrypsin, a cleavage product mass of 12,154.6 Da was identified, corresponding to residue Y64 in the switch II binding pocket as the cleavage site. CID fragmentation of the precursor ion m/z 608.8^+20^ confirmed this as aa65-aa169 C-terminal part of the protein (Figure 1C, S1, S2). The diminishing of these specific cleavage products in compound-bound samples may suggest that the binding of the compounds leads to conformational changes that protect this region from proteolysis, indicating the possibility that these residues are located within the switch II binding site. Indeed, the cleavage sites associated with compound binding were identified by LiP-MS at the switch II loop in accordance with crystal structure of the KRas G12D–Compound 5 complex (Figure 1A).

### Quantitative analysis of proteolysis extent

The degree of the proteolysis was measured quantitatively using the signal intensities of specific protein charge state ions at the maxima of their chromatographic peaks. For proteinase K, the signal intensity of the 698.95^+17^ m/z precursor was recorded, while for chymotrypsin, the signal intensity of the 715.99^+17^ m/z precursor was measured. The method is robust and rather fast (∼10 minutes per sample). The coefficient of variation (%CV) for the replicates was 7.12% for the proteinase K experiments and 9.97% for the chymotrypsin experiments, indicating high reproducibility of the measurements. Comparison across different compounds with varying affinities revealed differences in the degree of cleavage, correlating with the binding affinities and structural features of the compounds, with a notable exception of the Compound 2 (Figure 2). The method is also valid for the compounds with binding affinities in a micromolar range (data not shown). Quantitative measurements of the proteolysis reaction yields, their dependence on the location of the binding site, binding affinity and the particular compound structure makes this method potentially applicable for libraries screening to identify lead-like small molecules candidates.

**Figure 2.**
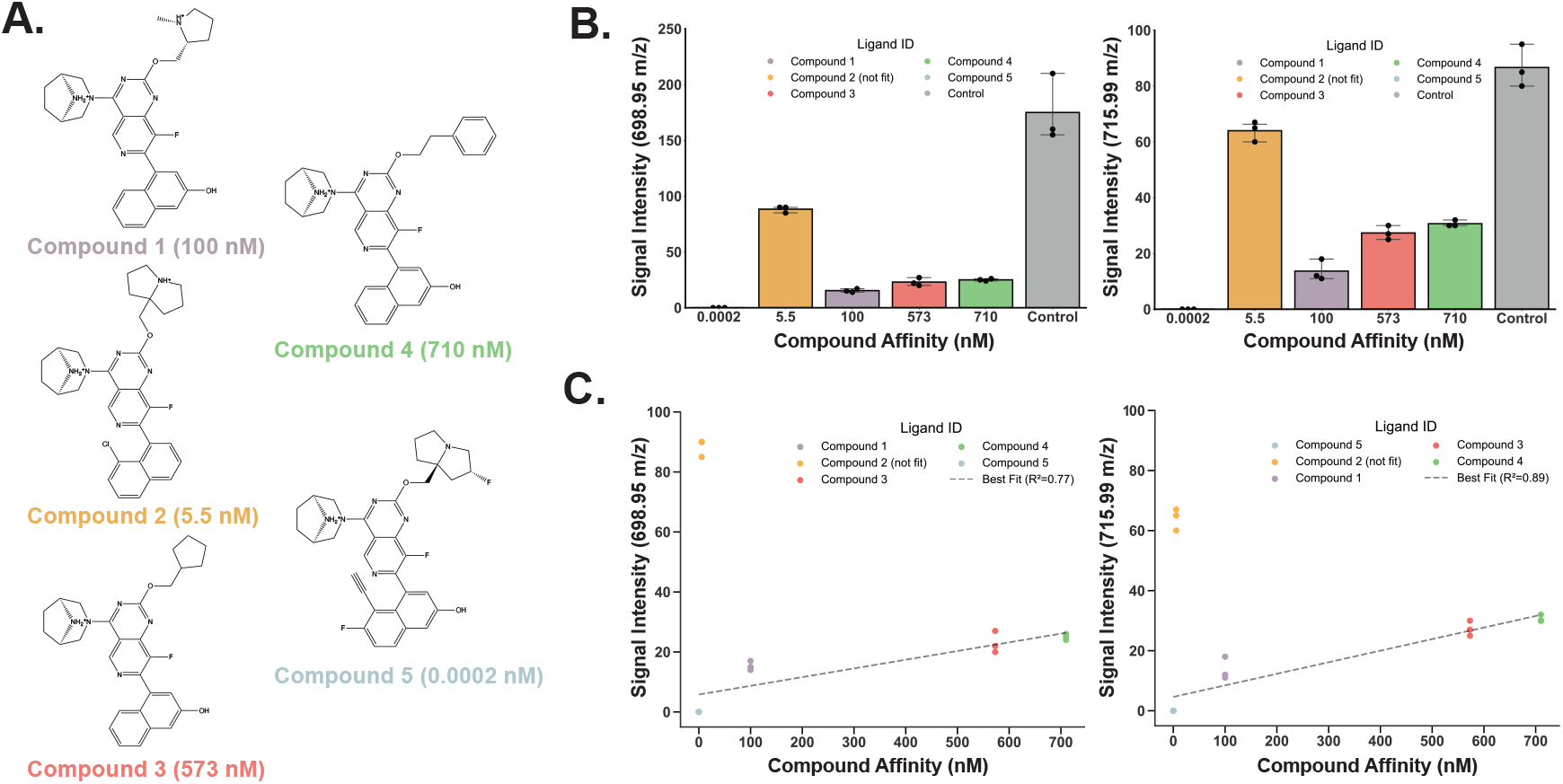
The degree of KRas G12D cleavage correlates with compound binding affinity. **A**. Structures of compounds used in this study and their KD values for KRas G12D. **B**. Degree of cleavage in KRas LiP-MS for compounds in this study. There are statistically significant differences in the degree of cleavage between the compound-bound and free states of KRas G12D for individual compounds. **C**. There is a correlation of the degree of cleavage in switch II loop and the KD of the ligands, with an exception of Compound 2 (Figure S3).

### Characterization of compound binding by MD simulations

Analysis of the MD trajectories for both the free and ligand-bound KRas G12D protein states revealed changes in the dynamics of the switch II region of the protein upon ligand binding. We found that the observed root-mean-square fluctuation (RMSF) values, which characterize relative flexibility of the protein along the entire peptide chain, get significantly reduced for the switch II region upon inhibitor binding when compared to the unbound protein complex (Figure 3A, 3B). Generally, proteases preferably cleave flexible locally disordered regions of the proteins (8). Upon inhibitor binding, stabilization of the switch II residues restricts the movement of the loop away from the protein core, resulting in reduced exposure of the cleavage sites to proteases (Figure 3C).

**Figure 3.**
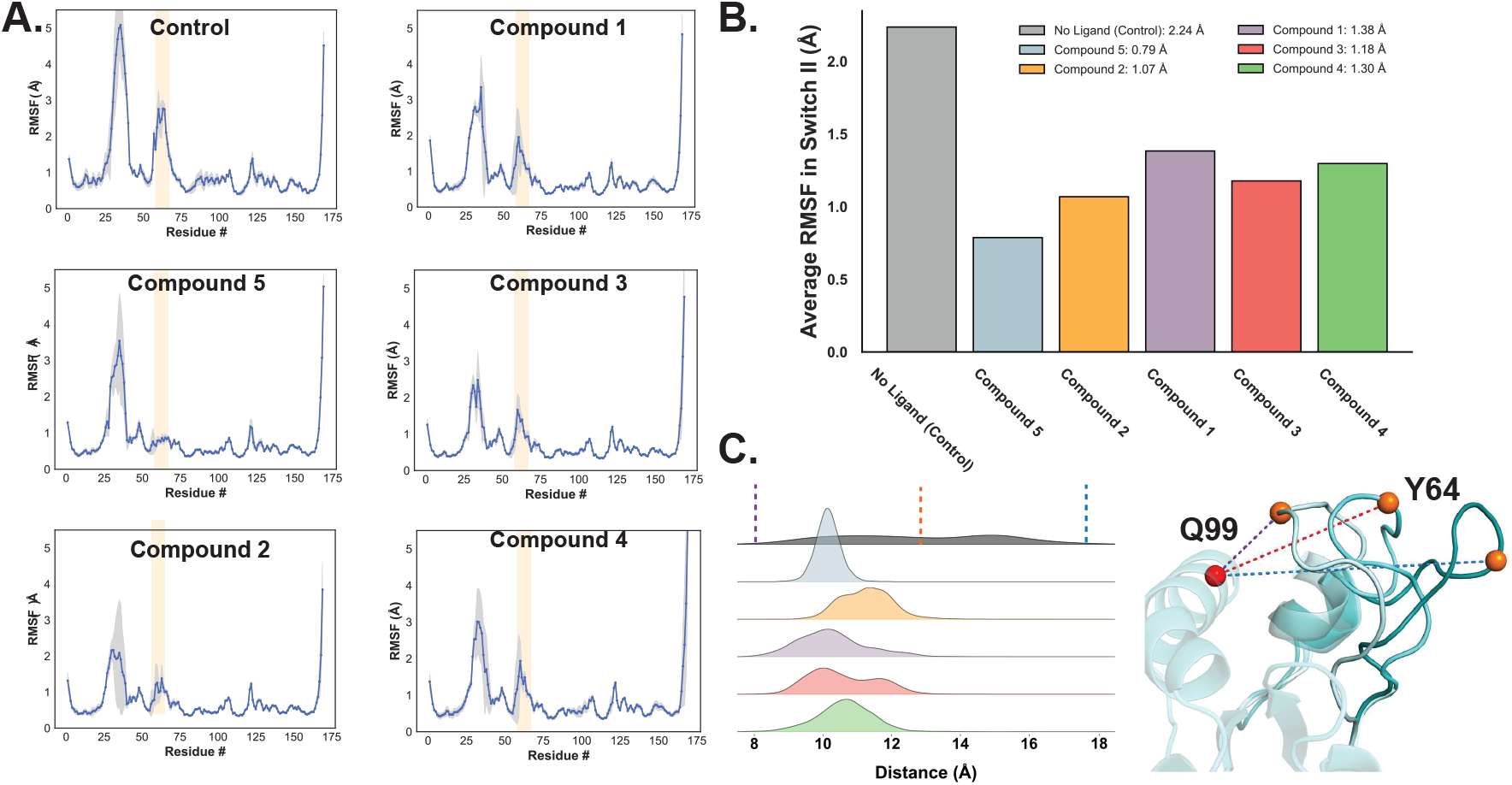
KRas G12D switch II region dynamics is modulated by compound binding. **A**. Changes in protein dynamics upon ligand binding, characterized by residue root-mean-squared fluctuation (RMSF). The average of three replicates is shown, with standard deviation indicated in gray. Ligand binding restricts the mobility of the switch II region (residues 58–67, highlighted in orange), with fluctuations decreasing as compound binding affinity increases. **B**. Average RMSF values for the entire switch II region, showing that higher affinity ligands result in lower average RMSF. **C**. Distribution of distances between cleavage site residue Y64 and reference residue Q99 (at the core of the protein). The width of the distributions reflects the magnitude of distance fluctuations during simulations. Compound binding and hydrogen bonding stabilize switch II fluctuations, as evidenced by the reduced distribution width, and pull the loop closer to the protein core, indicated by the leftward shift. Min (magenta), max (purple), and median (orange) distances are displayed as dashed lines on the distribution for unbound KRas. The same distances are shown on the corresponding snapshots from the MD simulation of the unbound KRas structure.

Higher-affinity ligands demonstrated greater conformational stability along the MD trajectories (Figures 4A, 4B), retaining more intermolecular interactions within the protein-ligand complex (Figures 4A, S4). This led to a more stable switch II region, corroborating the patterns observed in LiP-MS. The mobility of the ligand at the binding site modulates the mobility of the switch II loop and, consequently, increases the exposure of the cleavage sites to the proteases, which likely explains the observed correlation between binding affinity and the degree of proteolysis.

**Figure 4.**
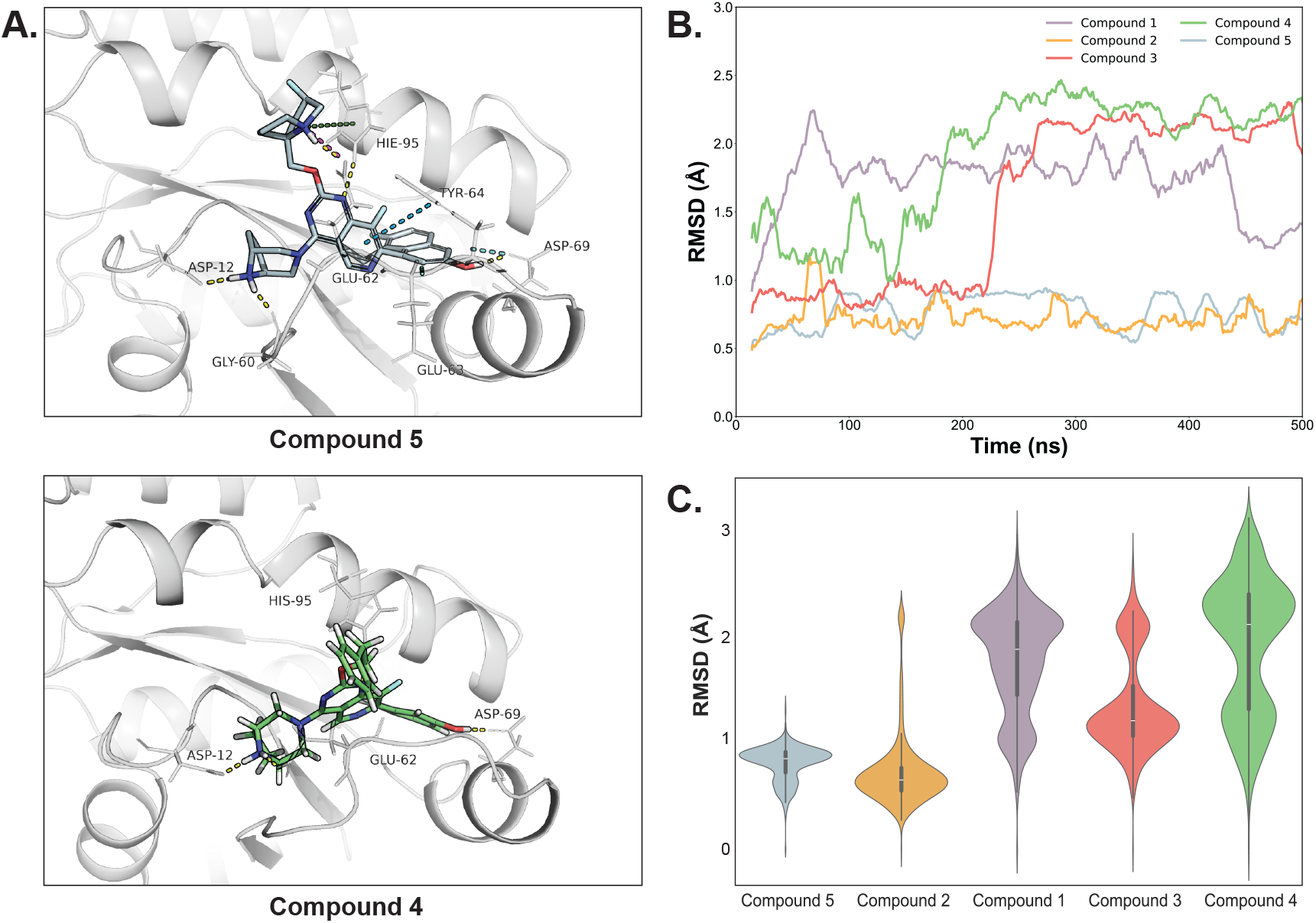
KRas G12D ligands with higher affinity exhibit greater stability and a more extensive interaction network. **A**. Snapshot of protein-ligand interactions with high affinity (Compound 5) and low affinity (Compound 4) ligands. The high-affinity ligand shows more interactions throughout the binding pocket (Figure S4) and forms a more extensive protein-ligand interaction network, including interactions with the switch II region. This correlates with the lower degree of proteolysis observed in our LiP-MS experiments. **B**. Dynamics of ligands bound to the KRas G12D. Lower ligand mobility (lower RMSD) is observed for high affinity ligands. Moving average smoothing function was applied for cleaner visualization purposes. **C**. Violin plots demonstrating the distribution of ligand RMSD values for each complex. Higher affinity ligands exhibit lower overall RMSD and reduced variation in RMSD values during simulations, indicating more stable ligand conformations.

In addition, the presence or absence of a particular functional group can also affect the extent of limited proteolysis. A detailed analysis of the KRas G12D – Compound 2 ligand simulation explained the deviation from the correlation trend between binding affinity and the degree of proteolysis observed for the other compounds. Despite its relatively high binding affinity, Compound 2 lacks an -OH group found in all other compounds, which plays a crucial role in coordinating interactions with residues Y64 and D69 through hydrogen bonding with the backbone nitrogen of Y64 and the terminal hydroxyl group of D69. The absence of stabilizing coordination at Y64 (Figure 3C) leads to greater exposure of the loop containing both proteolytic sites (Figures S3 and S4), compared to the other ligands in the series. This difference in protein-ligand interaction for Compound 2 explains both its high stability within the binding site and the increased flexibility near the cleavage sites, resulting in a higher degree of proteolysis. This approach, thus, provides a framework for selecting ligands that act through similar mechanisms of conformational stabilization, a critical feature for attenuating proper downstream signaling.

### LiP-MS-MD

Combination of quantitative LiP-MS and MD analyses provides insights into the mode of interaction of different compounds with KRas G12D and could be a valuable strategy for hit prioritization and lead optimization during hit-to-lead optimization. We show that LiP-MS can be used to identify compound binding sites on target proteins with high confidence and quantitative LiP-MS also provides insight into the degree of the protein structure stabilization upon compound binding for compounds within a chemical series. MD provides critical mechanistic interpretation of observed experimental proteolysis data for both protein structure stability changes and role of individual functional groups of the protein and the small molecule compounds in forming of the ligand-protein interaction network. Taken together, this information can be used for the intelligent selection of the lead compounds and further hit to lead optimization of chemical scaffolds (Figure 5).

**Figure 5.**
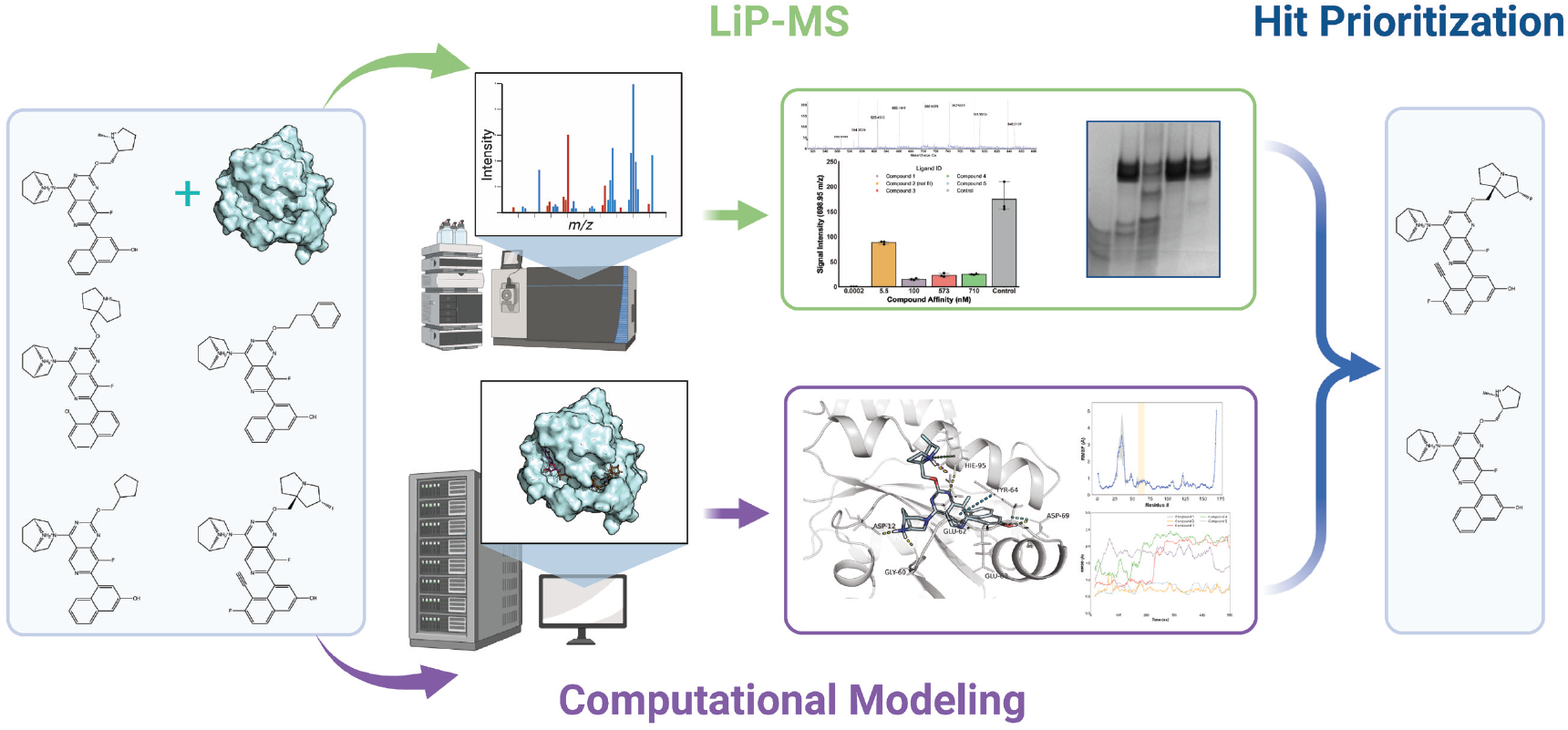
LiP-MS-MD. Parallel workflow for LiP-MS and in-silico analysis of KRas dynamics with ligands of varying affinities. Protein-ligand complexes are subjected to limited proteolysis and analyzed by LC-MS, with the degree of cleavage used to estimate protein structure stabilization. Docking studies are performed using Schrödinger’s Glide software, and all-atom molecular dynamics simulations are carried out using GROMACS with the CHARMM force field.

## Materials and Methods

### Protein and compound samples

Recombinant KRas G12D C118S (residues 1-169) protein was expressed in *E*.*coli* BL21 (DE3) RIL cells using a pET28a vector and purified to homogeneity using standard nickel affinity and size exclusion chromatographies. The C118S mutation has no effect on KRas G12D structure, but enhances the stability of Ras proteins (40). 10 mM DMSO stock solutions of compounds used in this study were stored until use at -20°C.

### Proteolytic enzymes and limited proteolysis reactions

For the limited proteolysis reactions, we used two proteolytic enzymes, chymotrypsin (TLCK treated, cat.# LS001432, Worthington) and proteinase K (recombinant, PCR grade, Roche Diagnostics GmbH). All the following solutions were made on the same day of the experiment. The stock solution of the Proteinase K was prepared in 50 mM acetic acid to a final concentration of 0.25 mg/mL. Chymotrypsin stock solution was prepared in LC-MS grade water to a final concentration of 0.1 mg/mL. KRas G12D stock solution was diluted in PBS (pH 7.4) to the final concentration of 10 µM. The compound solutions were each diluted in DMSO to the final concentration of 1 mM. 10 µL aliquots of KRas G12D protein solution (final concentration of 10 µM) were incubated with 1 µL of each of the compound solutions separately (final concentration of 100 µM, 1:10 protein-to-compound) in Eppendorf tubes at room temperature (23°C) for 1 hour. Three reaction replicates were prepared for each compound. Control samples were prepared by incubating 10 µL of the KRas G12D solution with 1 µL of DMSO under the same conditions. The solutions were then transferred to autosampler vial inserts. Each replicate and control sample was then incubated with 1 µL of the proteinase K 0.25 mg/mL at room temperature (23°C) for 5 minutes and subjected to LC-MS analysis. The same number of replicates and procedures were done using 0.1 mg/mL chymotrypsin as the proteolytic enzyme solution.

### SDS-PAGE

For SDS-PAGE analysis, the KRas G12D stock solution was diluted in PBS to a final concentration of 10 µM, and 30 µL of this solution was incubated with 3 µL of the Compound 5 solution (final concentration of 100 µM, 1:10 protein-to-compound) in 1.5 mL Eppendorf tubes at room temperature (23°C) for 1 hour. The control sample was prepared by incubating 30 µL of the KRas G12D solution with 3 µL of DMSO under the same conditions. The samples with Compound 5 and the control samples were then incubated with 3 µL of proteinase K (0.25 mg/mL) or chymotrypsin (0.1 mg/mL) at room temperature for 5 minutes. Following incubation, the reactions were quenched by addition of the 10 µL of the LDS sample buffer (Thermo), and the samples were loaded for SDS-PAGE analysis onto a 1 mm thick Bolt™ 4-12% Bis-Tris Plus precast gel (Thermo) along with molecular weight markers (Bio-Rad). The electrophoresis was performed at 120 V for 30 minutes, and the gel was subsequently fixed in 10% acetic acid/25% ethanol and stained with 0.01% Coomassie Brilliant Blue G-250 in 10% acetic acid/5% ethanol, followed by destaining in 10% acetic acid.

### LC-MS analysis

Following incubation, autosampler vial inserts containing proteolysis reaction samples were transferred to the autosampler and immediately injected for subsequent LC-MS analysis. Intact protein LC-MS analysis was performed using a Sciex TripleTOF 6600+ mass spectrometer coupled with the Shimadzu Nexera UPLC. AdvanceBio RP-mAb C4 column (2.7 µm particle size, 2.1 mm inner diameter × 50 mm length) was used. The analysis was performed using a binary gradient of (mobile phase A, 0.1% aqueous formic acid (FA); mobile phase B, acetonitrile containing 0.1% FA (v/v)) 0% to 100% B over 3 minutes, 100% B to 0% B in 0.1 min and maintaining 0% B for an additional 2.9 minutes at a flow rate of 200 uL/min. 1 ul injection volume was used. Total run time was 6 minutes. Full MS scans were acquired from 100 to 1500 m/z. The data were processed using Analyst 1.7.2 and PeakView 2.2 with BioTool Kit software (Sciex). For top-down analysis Information Dependent Acquisition (IDA) was used with an inclusion list of pre-selected precursor masses. The top 4 most intense precursor ions were isolated and fragmented with Collision-Induced Dissociation (CID) using a collision energy (CE) of 40 V. Precursor ions (792.0^+15^ m/z for Proteinase K cleavage product and 608.8^+20^ m/z for chymotrypsin cleavage product) were included for MS/MS analysis at a retention time of 3.6 minutes for 60 seconds, with an intensity threshold of 1 cps. The cleavage product masses were matched to the theoretical masses of all possible peptides of the KRas G12D protein. Detected fragment masses were searched using the Protein Prospector website (https://prospector.ucsf.edu/prospector/mshome.htm).

### Quantitative measurements of the proteolysis extent with different compounds

To quantitatively assess the degree of cleavage among different compound samples with varying affinities and structures, the signal intensities of specific precursors at the maxima of the corresponding chromatographic peaks were measured and compared. For proteinase K, the signal intensity of the 698.95^+17^ m/z peak was recorded, while for chymotrypsin, the signal intensity of the 715.99^+17^ m/z peak was measured.

### Structure preparation and docking

All *in silico* experiments were conducted using the nucleotide-bound form of KRas G12D mutant in complex with MRTX1133, PDB ID: 7RPZ. Prior to docking, the structure of the protein was prepared using Schrödinger Protein preparation tools with default settings: adding and optimizing hydrogen atoms, setting residue ionization and tautomer states with PROPKA at pH 7, removing water molecules more than 3 Å away from the HET atoms, and performing restrained structure minimization with the OPLS4 force field to converge the heavy atoms to an RMSD of 0.3 Å (41, 42). This offers minor structure optimization to ensure optimal performance of subsequent docking runs. Following protein preparation, the Glide receptor grid was generated, with the grid box centered on 6IC ligand and inner and outer box sizes set to 10 Å and 26.75 Å, respectively. The structures of all ligands were prepared using the LigPrep tool to sample all possible conformation states during the docking protocol (43). Docking was then conducted for the series of ligands prepared on the previous step. The best models were selected based on the Glide SP docking score, with the top-scored pose chosen as the structural model of the complex (44). These structures were then used as inputs for downstream molecular dynamics simulations.

### Molecular dynamics simulations

The protein-ligand complex models identified through docking were used as initial structures for MD simulations using GROMACS software. All ligands (compounds and GDP) were individually prepared in PyMOL, then parametrized for CHARMM36 force fields using SwissParam and CHARMM-GUI (45-48). The unit cell was established as a rhombic dodecahedron with the protein >= 1 nm from all edges. The protein structure was solvated using the TIP3P water model and neutralized with sodium and chloride ions. Particle mesh Ewald (PME) was employed for long-range electrostatic interactions, with a 10 Å cutoff for non-bonded interactions. The system was initially equilibrated using an NVT ensemble, followed by further equilibration under the NPT conditions. GDP and compounds were individually restrained during minimization and equilibration. Production runs were conducted using GROMACS 2020.3 on UNC high-performance computing clusters with Nvidia V100 GPUs. Simulations were at a constant pressure and temperature of 1 atm and 270 K for 500 ns in three replicates with identical parameters. All replicates were individually minimized and equilibrated to obtain unique initial velocity distributions (49, 50).

### MD data analysis

MD trajectories were processed using the MDTraj package in Python. Briefly, trajectories were exported from GROMACS after removal of the periodic boundary condition and centering of the protein in the simulation cell. All simulations were exported with a timestep of 1 ns for a total of 500 frames. Analysis was then conducted by selecting protein and ligand structures programmatically and analyzed over the course of the entire 500 ns trajectory. In our analysis, the overall dynamics and flexibility of the complex were characterized by the backbone carbon atom root-mean-square fluctuations (RMSF) for the protein and by root-mean-square deviation (RMSD) for the small molecule ligands. To account for the extensive ligand fluctuations during the simulations, we applied a moving average smoothing function to better visualize their dynamics. The algorithm processes the time series data by averaging each point with its neighboring frames, creating a smoothed value for each time point to enhance trend clarity (51). Structural visualizations were generated with PyMOL. Protein-ligand interaction fingerprints were obtained using the ProLIF python package to isolate ligand snapshots and determine intermolecular interactions occurring with a timestep of 10 nanoseconds for a total of 50 snapshots (52).

## Acknowledgments

This work was supported by funding to CHB for “The Metabolomics Innovation Centre” from Genome Canada through the Genomics Technology Platform (265MET and MC4T), and by grant #42495 to CHB from the Canadian Foundation for Innovation via the Major Sciences Initiatives Fund. CHB is also grateful for support from the Segal McGill Chair in Molecular Oncology at McGill University, and for support for the Segal Cancer Proteomics Centre at the Jewish General Hospital (Montreal, Quebec, Canada) from the Warren Y. Soper Charitable Trust and the Alvin Segal Family Foundation. The authors gratefully acknowledge support from the NIH Biophysics Training Grant (T32GM148376-01A1).

