## Supplemental Material for "A Hit Prioritization Strategy for Compound Library Screening Using LiP-MS and Molecular Dynamics Simulations Applied to KRas G12D Inhibitors"

**Supplementary data.**

Figure S1. Limited proteolysis of KRas G12D in free and compound-bound states.

Figure S2. Top-down analysis of the KRas G12D cleavage products.

Figure S3. LiP-MS pattern correlates with ligand stability and coordination of cleavage site residues.

Figure S4. Protein-ligand interaction networks show more extensive and stable interaction patterns for high-affinity ligands.

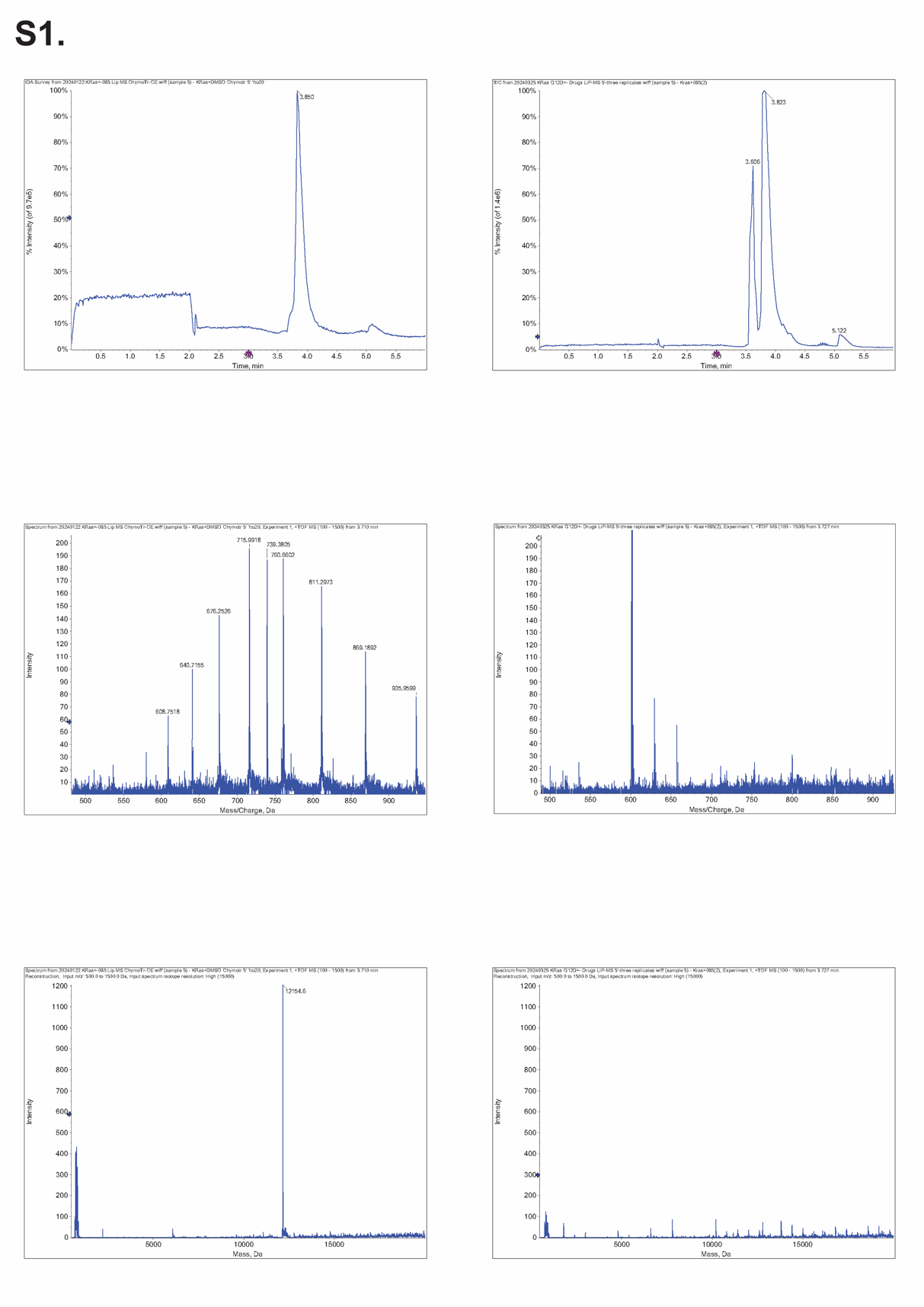

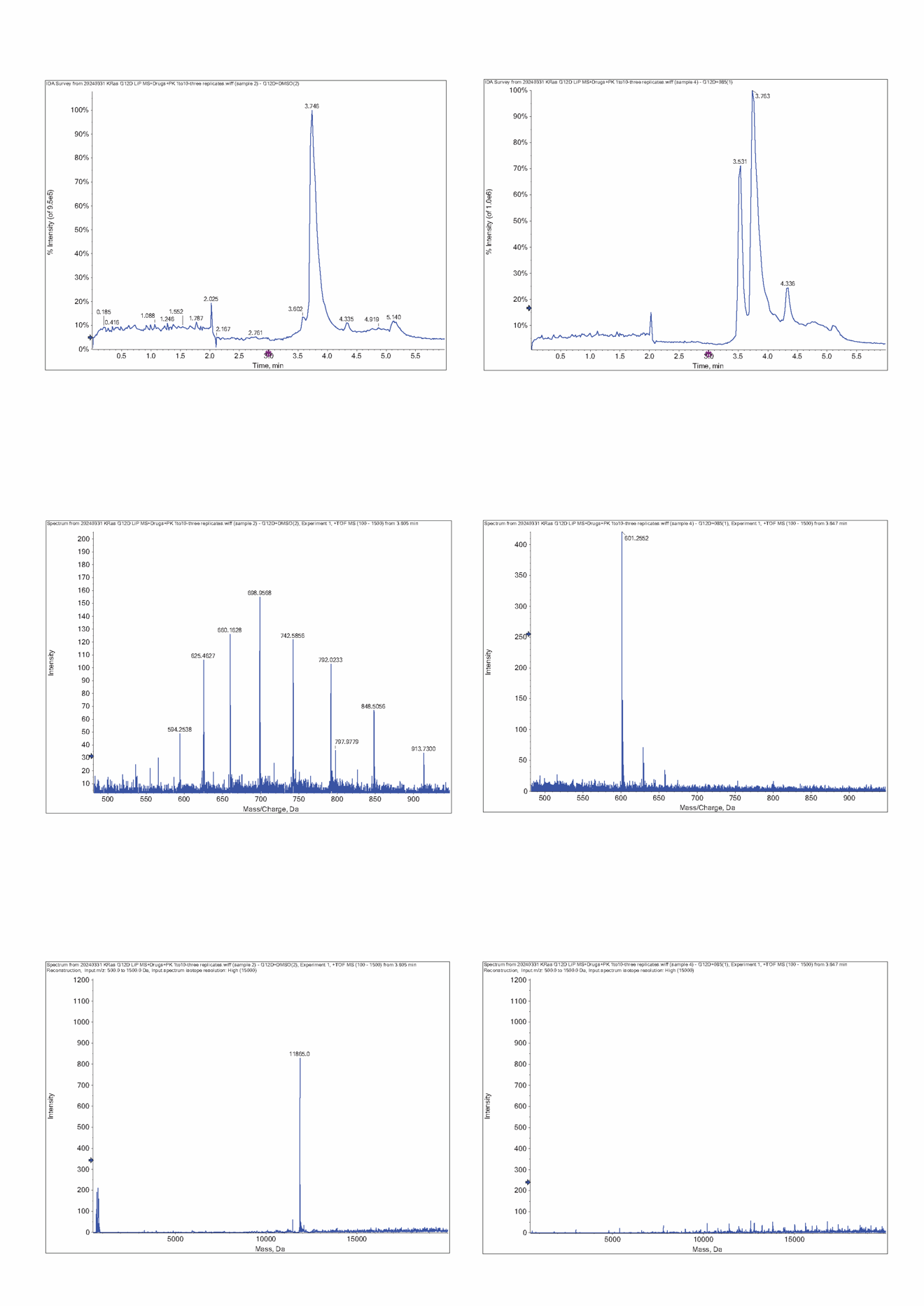

**Figure S1. Limited proteolysis of KRas G12D in free and compound-bound states.** Intact protein LC-MS analysis of KRas G12D limited proteolysis reactions using chymotrypsin (top) and proteinase K (bottom) without (left) and with (right) Compound 5. From top to bottom - total ion chromatograms, representative charge envelopes and the corresponding deconvoluted mass spectra of the protein cleavage products. There is a noticeable absence of the protein cleavage products in the ligand-bound samples.

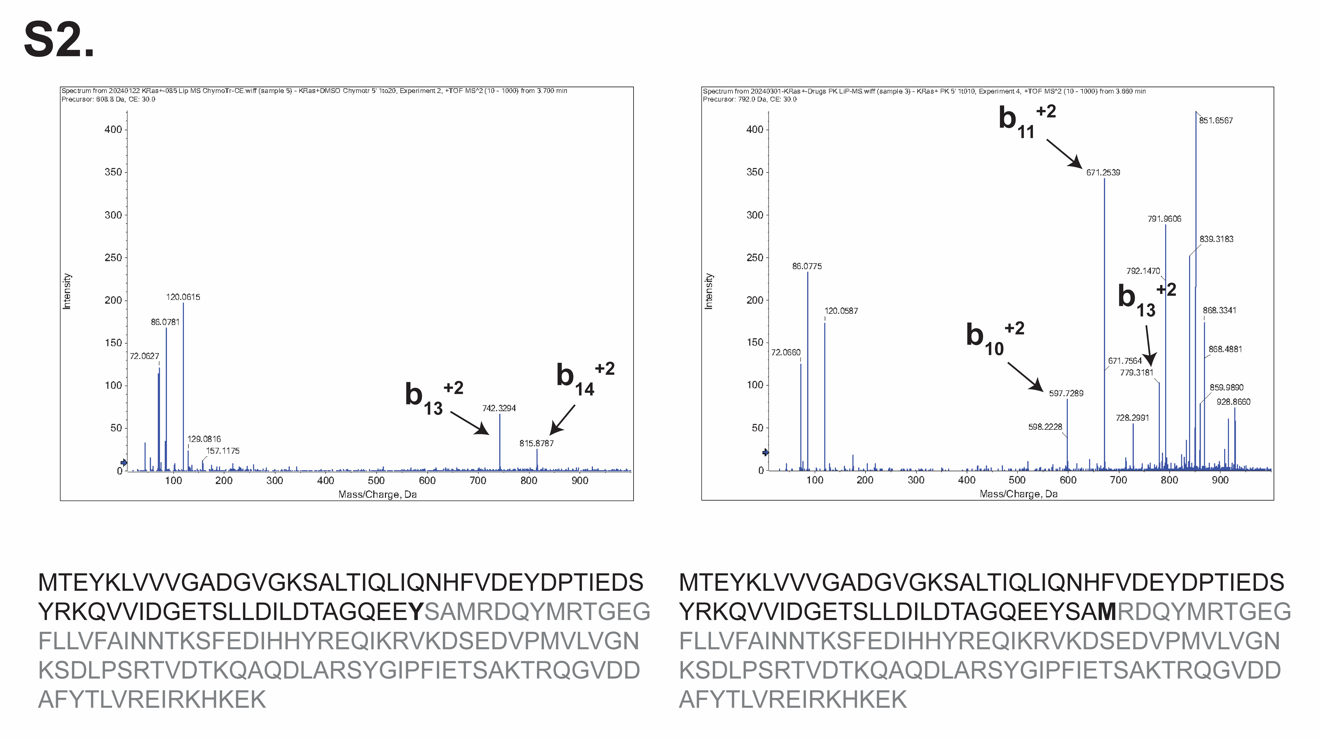

**Figure S2.** **Top-down analysis of the KRas G12D cleavage products.** Left, MS/MS mass spectrum of the KRas G12D chymotrypsin cleavage product with 12154.6 Da mass (608.8^+20^ m/z precursor). Right, MS/MS mass spectrum of the KRas G12D proteinase K cleavage product with 11865.0 Da mass (792.0^+15^ m/z precursor). The cleavage products were identified as C-terminal parts of the KRas G12D protein with the cleavage sites Y64 and M67 for chymotrypsin and proteinase K, respectively. These sites are highlighted on the KRas G12D sequence.

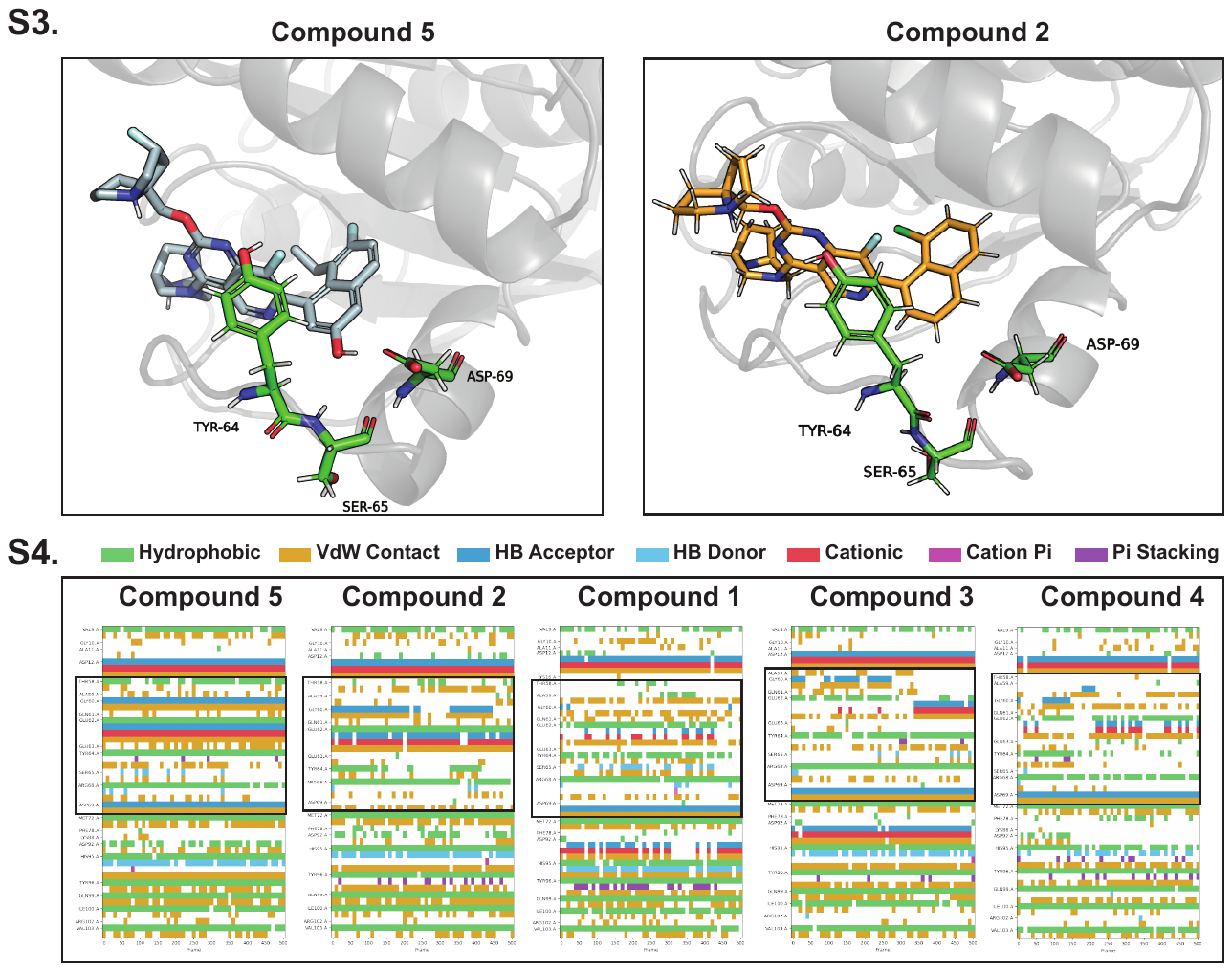

**Figure S3. LiP-MS pattern correlates with ligand stability and coordination of cleavage site residues.** Simulation snapshots of Compound 5 and Compound 2 show the interaction of the -OH group of Compound 5 with residues Y64 and D69. This interaction is absent in Compound 2. Additionally, minimal transient hydrogen bonding occurs between the -OH group and residue S65 during the simulation (Figure S4). The absence of these interactions may explain the deviation in the correlation between binding affinity and degree of cleavage in the switch II loop for Compound 2.

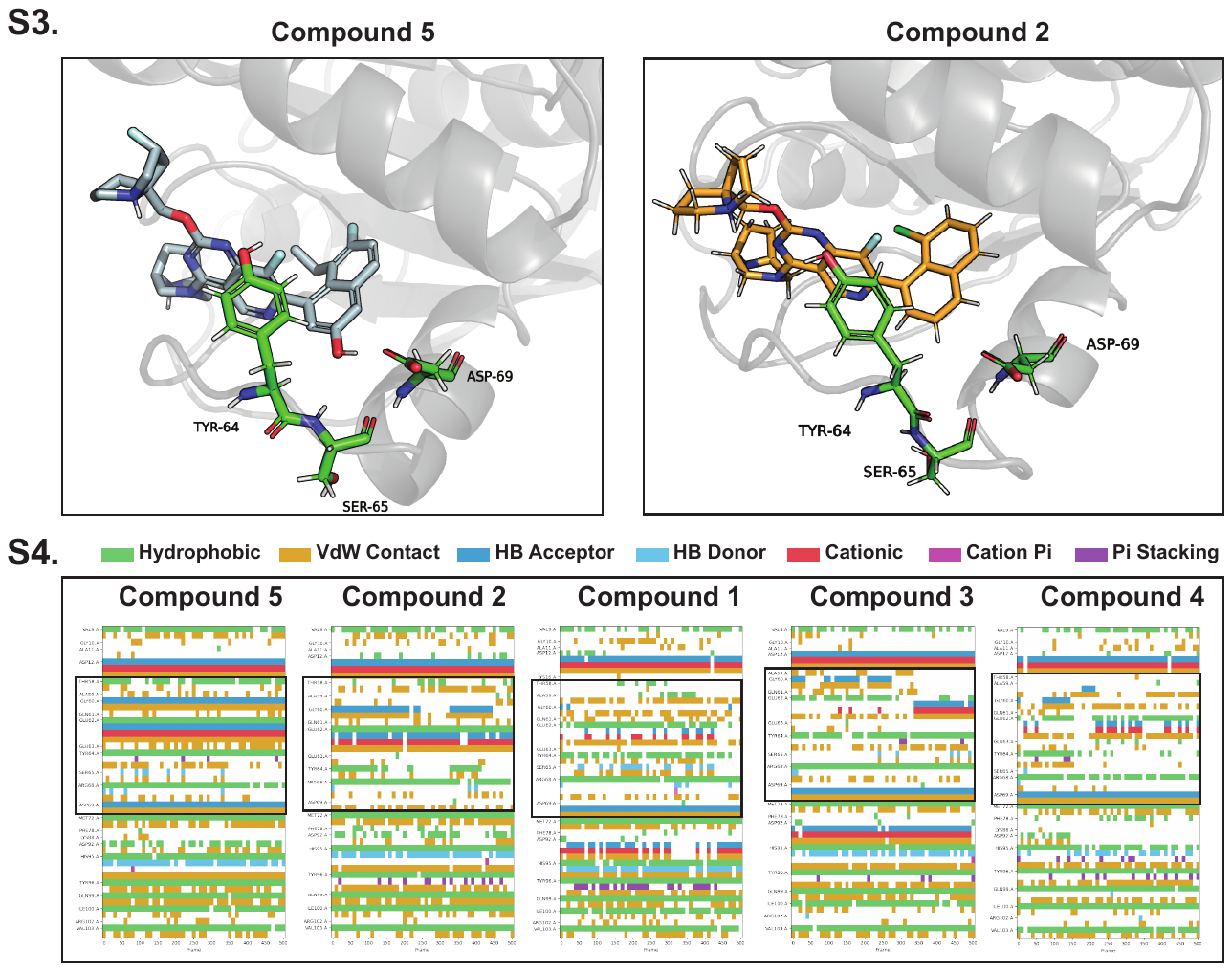

**Figure S4. Protein-ligand interaction networks show more extensive and stable interaction patterns for high-affinity ligands.** Interactions between KRas G12D and the ligands were computed over the course of the simulations. The black box highlights the switch II region for each complex. Higher-affinity ligands exhibit more interactions, with a more consistent pattern throughout the simulation, contributing to the stabilization of both the ligand and the surrounding protein pocket.
